# Leveraging synthetic environmental covariates to explore crop genotype by environment interactions

**DOI:** 10.64898/2026.10.15.759392

**Authors:** Stephanie M Brunner, Hans-Peter Piepho, Karine Chenu, Bita Heidariask, David Moody, Silvina Baraibar, Dini Ganesalingam, Lee Hickey, Kai P. Voss-Fels, Hannah Robinson

## Abstract

Environments are unpredictable. Genotype by environment interactions (GEI) further complicate any assessment of genotype performance or stability across diverse environments posing a challenge for plant breeders. Several approaches exist to address this challenge. However existing approaches present a trade-off between scale and interpretability often acting in a ‘black box’ manner. Here we extend the Finlay–Wilkinson regression on synthetic environmental covariates of Piepho and Blancon (2023) to dissect GEI at the level of individual developmental windows, resolving which growth stages drive GEI and how genotype stability varies across them. Yield data for 961 genotypes from a barley breeding program, tested across 12 environments, is used as a case study to illustrate how this approach can provide valuable insight into the GEI present. Exploring the dynamic impact of environmental variables across growth stages revealed that environmental factors during early grain fill contributed the most to yield GEI variance. Stability was found to be dynamic across development windows, presenting growth-stage-specific yield stability as a new potential breeding target. This approach yields new, growth-stage-resolved measures of yield stability and a new means of dissecting GEI in complex traits, while also opening a route to environmental clustering based on stage-specific GEI patterns.

**Highlight:** Application of synthetic covariate modelling can reveal drivers of genotype by environment interactions and provide a new understanding of stability for crop breeding.

## 1 Introduction

Environmental conditions have always been unpredictable. Even with the constant improvement of forecasting models there is a trade-off between lead time and accuracy which can seriously impact agricultural decision making (Meza *et al*., 2008). Climate change intensifies this challenge by increasing the frequency of extreme events such as droughts and flooding (Newman and Noy, 2023; Heidariask *et al*., 2026). Out-of-season temperature changes, particularly at sensitive times developmental stages, can also reduce yield (Zheng *et al*., 2012; Hill *et al*., 2024; Ullah *et al*., 2024). Recent modelling indicates that climate change reduced wheat yield potential by up to 12% between 1990 and 2023, in particular due to heatwaves (Ababaei and Chenu, 2020; Lawes *et al*., 2025). Understanding how environmental conditions at key developmental stages drive yield losses is therefore critical. This is further complicated by genotype-by-environment interactions (GEI), whereby genotypes respond differently to environmental conditions.

Crop breeders select for high performance across a target population of environments (TPE), aiming to maximise genetic gain under conditions relevant to commercial production (Heidariask *et al*., 2026). A well-characterised TPE should be strongly correlated with the target production environment so that genetic gain translates beyond the breeding program (Cooper et al., 2023). However, TPEs have often been defined by using geographic proximity or micro-climatic zones rather than environmental factors that drive GEI (Chenu, 2015).

Multi-environment trial analyses show that year can contribute more to GEI than location, yet year-to-year effects are substantially harder to predict and interpret (Smith *et al*., 2001, 2015, 2021; Bustos-Korts *et al*., 2022).

Much of our mechanistic understanding of how specific environmental factors, such as temperature, influence key traits comes from controlled glasshouse experiments (Ochagavía *et al*., 2022; Slawin *et al*., 2024). While these studies are valuable for establishing the fundamental basis of such responses, highly controlled conditions rarely translate to large-scale field trials and commercial breeding environments (Poorter *et al*., 2016). This leaves a substantial gap in understanding which environmental factors drive GEI under real-world conditions, and how they influence key trait performance. Furthermore, with growing season or “year” increasingly recognised as the dominant source of GEI, there is a growing need to understand broader environmental trends and their interactions with specific genomic regions across the growing season.

One approach that breeders have taken to manage GEI is to select for stability, by identifying genotypes that perform consistently across a range of environments. However, breeding for stability carries trade-offs. Evidence suggests that the most stable genotypes are not necessarily the highest-performing in any individual environment, meaning breeders may sacrifice yield potential in pursuit of stability (Sinebo, 2005; Ndiaye *et al*., 2019). Despite these limitations, stability selection has remained a practical default in many breeding programs, largely because the methodological tools needed to dissect GEI at scale have not been readily available. Without scalable approaches to characterise GEI, selecting stable and high yielding varieties remains imprecise, and the potential to exploit genotype-specific environmental responses goes largely unrealised.

From the early days of recognising these GEI, there have been a range of approaches by which to model them including the Finlay-Wilkinson regression, Additive Main Effects and Multiplicative Interaction modelling and various visualisation tools such as biplots which allowed breeders to understand which varieties had the most stable performance across the tested environments (Finlay and Wilkinson, 1963; Zobel *et al*., 1988; Rahmati *et al*., 2024). With the advancement of statistical modelling have come more complex and complete ways to model GEI data, which provides more insight into the relationship between the genotype and the environments rather than simply a ranking. One such approach, widely used by breeding programs, is the factor-analytic (FA) model (Piepho, 1998; Smith *et al*., 2001). This purely latent modelling approach defines a series of factors which account for the main (first factor) and subsequent variance contributing to the GEI (Smith *et al*., 2019). This approach works effectively in prediction-based models, such as GBLUP, to capture the GEI across large numbers of field trials and can be used to identify the most stable varieties across those environments (Smith *et al*., 2021; Piepho and Williams, 2024). FA models excel at capturing large amounts of GEI by defining latent factors from variance structures in the data.

However, their purely latent nature precludes formal testing of specific environmental drivers on the GEI patterns identified (Tolhurst *et al*., 2022). While environmental covariates have been correlated with FA factors post-hoc in some datasets, this approach is not ideal, as environmental data is not formally incorporated into the modelling framework (Fradgley *et al*., 2025). Compounding this, the common practice of fitting environment as a fixed effect in FA models restricts interpretability and limits prediction to only those environments included in the dataset (Piepho and Williams, 2024). In contrast a reaction norm approach can be performed across a single environmental axis allowing for clear interpretation and visualisations of the trends across environments for specific genotypes (Ly *et al*., 2017, 2018; Waters *et al*., 2023). However, extending this framework to incorporate additional environmental variables as individual random interaction terms rapidly becomes computationally unfeasible (Jarquin *et al*., 2014; Piepho and Blancon, 2023). More complex random regression approaches can accommodate multiple environmental covariates through an environmental relationship matrix (Jarquin *et al*., 2014), but in doing so lose the ability to directly model individual genotype-by-environment covariate interactions, resulting in a subsequent loss of interpretability.

An approach to facilitate interpretation whilst asking broader questions on the nature of GEI is a combination of the latent component of the FA model integrated with known covariates through random regression as outlined in Tolhurst et al. (2022). Combining effective pieces from both FA model and reaction norms allows for more genetic variance to be captured by such a model and for increased understanding on the relationship between environmental variables and the trait of interest (Tolhurst *et al*., 2022). However, there are limitations in the ability to predict into future environments with this approach as the FA model component requires a fixed environmental effect (Piepho and Williams, 2024). Further, the genotype sensitivities are also not made explicit in this approach, as they are modelled jointly with the intercepts within the factor-analytic structure and remain confounded with them (Piepho and Blancon, 2023). An alternative approach which allows for the fitting of a single-stage GBLUP model is the Extended Finlay-Wilkinson regression model with environmental covariates as described by Piepho and Blancon (2023). Returning to the fundamentals of Finlay-Wilkinson regression, this approach allows individual environmental covariates to be fitted directly within the model, retaining the interpretability of genotype-specific sensitivity parameters analogous to the slopes in classical Finlay-Wilkinson regression. To address the challenge of high-dimensional environmental data, the approach introduces the concept of synthetic covariates, which are linear combinations of observed environmental covariates that reduce dimensionality whilst retaining the ability to capture substantial GEI variance (Piepho and Blancon, 2023). Importantly, this allows a greater number of environmental covariates to be accommodated within a parsimonious model without requiring a latent component, thereby maintaining a direct link between model parameters and observable environmental variables. Several methods for deriving synthetic covariates are described by Piepho and Blancon (2023), including criss-cross regression, partial least squares, redundancy analysis, and a factor-analytic approach, each offering different benefits and trade-offs. A key advantage of this framework is that, once synthetic covariates are derived, predictions can be extended to new environments not represented in the training dataset, provided environmental covariate data is available for those environments. Additionally, synthetic covariates are weighted by their contribution to GEI, meaning that environmental covariates are not treated as having equal influence on genotypic performance in the final model.

The current body of work utilising this approach focuses primarily on prediction into new environments using the environmental covariates (Tadese *et al*., 2024, 2025; Hrachov *et al*., 2026). However, though speculated, the genomic relationship matrix (GRM) is yet to be incorporated into the framework as is common practice in exploring breeding program populations. Additionally, this approach has the potential to be expanded to help represent crop physiology and response to environment on a large scale to dissect the nature of GEI. The combination of these, as will be shown in this paper, can allow for the exploration of environment by haplotype interactions identifying target regions for breeders for stability or specificity. The simplistic complexity of this model provides a playground for understanding the drivers of GEI beyond its intended purpose - to maximise the GEI captured - to a tool that can be leveraged to explore complex questions in a multi-environment data set.

This paper develops a pipeline to explore the environmental sources of GEI in a multi-environment data set of a barley breeding program, utilising the extended Finlay-Wilkinson regression on synthetic environmental covariates. Leveraging real world breeding program data on barley yield across multiple years in Australia a flexible and interpretable pathway to dissect the environment component of GEI will be explored. The aims of this paper are: 1) To develop a framework for understanding the environmental drivers of GEI at specific developmental stages expanding on the Extended Finlay-Wilkinson regression approach. 2) To apply this framework in a barley breeding program data set to illustrate its potential application in commercial breeding programs. 3) To provide examples and discussion regarding how this pipeline can be applied broadly and provide valuable information for selection decisions.

## 2 Materials and Methods

### 2.1 Datasets

#### 2.1.1 Phenotype and genotype data

961 barley genotypes were assessed for yield performance in 12 field trials across six locations and four years (Supplementary Table 1). Germplasm data was provided by Australian breeding company InterGrain Pty Ltd. Genotype data was obtained using wheat barley 40K v1.9 BeadChip returning 12,548 high quality single nucleotide polymorphism (SNPs) for barley (Keeble-Gagnere *et al*., 2021). Markers were curated to ensure all remaining markers had <10% heterozygosity and >0.05 minor allele frequency resulting in 6599 final SNPs. Principal component analysis was used to explore the genetic structure present in the data set. A genomic relationship matrix (GRM) was then generated using R package ‘AGHmatrix’ and the Van Raden method (Amadeu *et al*., 2023). This data set has been previously described in Brunner *et al*. (2026).

#### 2.1.2 Weather data

Weather data was obtained from the SILO based data set which provides interpolated data for daily weather across Australia based on collated weather station data (Jeffrey *et al*., 2001).

Weather data was extracted for specific trial dates and locations with R package ‘EC4MET’ which also provided the calculation for soil moisture index (SMI) (Fradgley and Whan, 2025; Fradgley *et al*., 2025).

#### 2.1.3 Phenology data and modelling

To align weather data respective to phenology, flowering time information was required for all trials. Zadoks scores or days to flowering was collected at all trials except YRK21 (Supplementary Table 1). To predict flowering date in trials without days to flowering, trials were simulated for the cultivar Spartacus CL using APSIM-Barley crop model (version 2024.6.7514.0) (Holzworth *et al*., 2014, 2018). Simulations were driven by daily weather data obtained from the SILO based dataset, soil properties from a nearby characterised profile in the APSoil database (Jeffrey *et al*., 2001; Dalgliesh *et al*., 2012), and trial-specific management practices. As no initial soil water measurements were taken, simulations were initiated six months prior to sowing to capture pre-season environmental conditions and allow soil water recovery, with initial plant available water (PAW) set to 20% and uniformly distributed throughout the soil profile. To best simulate crop growth and development at each trial, the cultivar-specific parameter ‘minLN’ (minimum leaf number under non-limiting vernalisation and long photoperiod) was finetuned so that simulated phenology aligned with field-observed Zadoks scores, where recorded (Christopher *et al*., 2016; Amin *et al*., 2025).

To ensure consistency across trials and to obtain unbiased, spatially adjusted estimates of days to flowering, in trials where the data was collected, flowering time for Spartacus CL was estimated as best linear unbiased estimators (BLUEs) from a spatial mixed model:

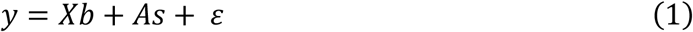

in which *y* is the response vector for days to flowering for each variety in each trial, *b* is a vector of fixed effects which includes genotype and fixed effects associated with trial design (i.e. column and row) when statistically significant, *s* represents a vector of random effects from spatial patterns (i.e. column and row effects where significant), both *X* and *A* are design matrices. *ε* represents the plot level residual error modelled with an autoregressive AR(1) x AR(1) variance structure. The combined information of sowing and harvest dates were then all converted to growing degree days relative to Spartacus CL flowering.

### 2.2 Synthetic Environmental Covariate Pipeline

The synthetic environmental covariate pipeline consists of four key steps with two different mixed linear models as shown visually in Figure 1. Environment data is divided into biologically relevant groups (e.g. by developmental windows) such that in each group there is a single value for each environmental variable in each environment. The dimensionality of each group must be such that the number of environmental variables is less than the number of environments. The environmental covariate model is then fit to the data set (see 2.2.1), from this model the factor loadings are extracted and used to derive synthetic environmental covariates. The environmental covariate model is then fit multiple times once for each group until the desired number of synthetic environmental covariates have been derived. Finally, the synthetic covariate model is fit utilising the synthetic environmental covariates to capture the GEI.

**Figure 1.**
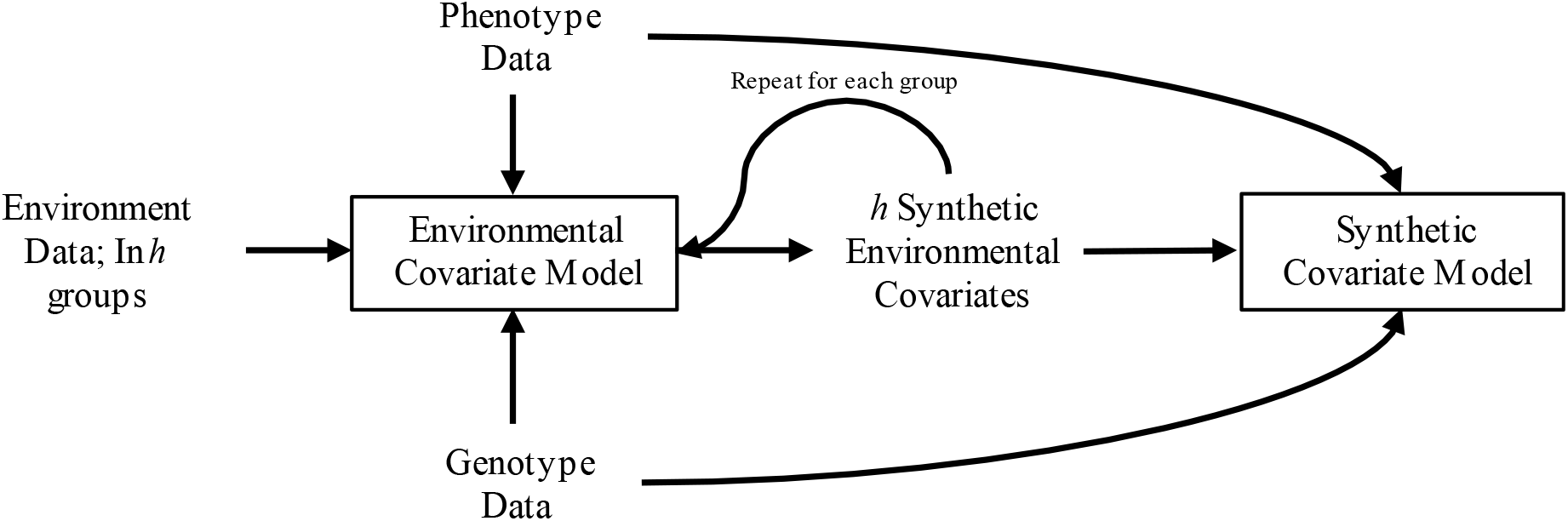
*Graphical representation of synthetic environmental covariate approach and its inputs. h* is the number of synthetic environmental covariates fitted in the synthetic covariate model. It equals the number of groups the environmental data are divided into, as a separate environmental covariate model is fitted to each group.

#### 2.2.1 Environmental covariate model

The environmental covariate model is an extended Finlay-Wilkinson regression model utilised to derive the synthetic environmental covariates (*z*_*j*_) with the same phenotype and genotype data that will be included in the final model (Figure 1). The environmental covariates are centred and scaled (z-standardised) prior to fitting the model. The model is fit in line with Piepho and Blancon (2023):

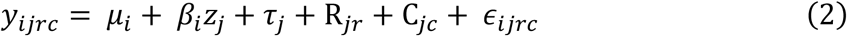

where *y*_*ijrc*_ is the observed plot level yield of genotype *i* (*i =* 1, …, *n*) in environment *j* (*j* = 1 ,…, *k*) and *μ*_*i*_is the fixed genotype effect representing the mean performance of genotype *i, β*_*i*_is the genotype-specific slope as calculated from the random regression interaction between the genotypic effect (modelled using a GRM) and the environmental covariates, *τ*_*j*_represents the random main environmental effect, R_*jr*_ and C_*jc*_ represent the environment specific random effects of row and column, and *z*_*j*_ is the synthetic environmental covariate for environment *j* derived from

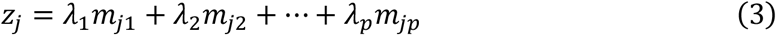

where *m* is the environmental covariate (*m* ∈ {1,2 … *p*}) and *λ* is the weighting associated with the first factor from a random regression model (*λ* ∈ {1,2 … *p*}) applied to the covariance between the genotype and environment terms. The only outputs of this model used in downstream analysis are the synthetic environmental covariates. This model is run multiple times (Figure 1) with each new group of environmental covariates to create a matrix of synthetic covariates with dimensions given by the number of environments (*k*) by the number of synthetic covariates (*h*).

#### 2.2.2. Variable and synthetic covariate definition

To understand the importance and changing impact of environmental factors on GEI across growth stages, synthetic covariate groupings were determined based on summed environmental values within developmental windows, centred at flowering of Spartacus CL. With limited environmental data available the seven environmental covariates (*m* ∈ {1,2 … 7}) were selected to be summed with in each time window: rainfall (mm), maximum temperature (°C), minimum temperature (°C), vapour pressure deficit (VPD) (hPa), radiation (MJ/m^2^), day length (minutes) and soil moisture index (SMI). Summed values were selected to represent cumulative environmental exposure rather than average conditions and were standardised before analysis.

To identify the developmental windows represented in this data set, 11 developmental windows, each spanning 100 or 200°Cd were defined across the thermal time span shared by all environments from 1100°Cd before Spartacus CL flowering to 1000°Cd after Spartacus CL flowering. The environmental covariate model (Eq. 2) was fit once for each developmental window and pairwise correlations between the associated *λ* were evaluated with Pearson correlation analysis. Chronologically adjacent windows with positive correlations between their *λ* were grouped together, resulting in six developmental windows. These windows are referred to hereafter as emergence, development, flowering, early grain fill, late grain fill and maturity. These windows do not represent the exact timepoint at which each physiological stage was reached, as only flowering time is explicitly measured or calculated in this study. However, the windows have been named given their relative position in relation to sowing, flowering and harvest. The environmental covariate model (Eq. 2) was then fit six times, once for each of the identified growth windows, to derive synthetic covariates at each (*z*_*jh*_with *h* ∈ {1,2 … 6}).

#### 2.2.3 Synthetic covariate model

The synthetic covariate model takes the six synthetic covariates and fits an extended Finlay-Wilkinson regression model (Figure 1). The model is fit as a modified version from Piepho and Blancon (2023):

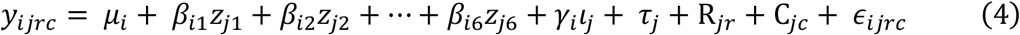

where *y*_*ijrc*_ is the observed plot level yield of genotype *i* in environment *j, μ*_*i*_ is the fixed genotype effect, *β*_*ih*_ is the genotype-specific slope, representing the sensitivity of genotype *i* to synthetic covariate for the growth window *h*, modelled through an unstructured variance-covariance matrix constrained by the GRM, *γ*_*i*_*ι*_*j*_ is the genotype-specific slope describing the interaction between the GRM and residual GEI not captured by the synthetic covariates, fitted with a diagonal variance-covariance structure assuming independent environments. R_*jr*_ and C_*jc*_ represent the environment specific random effects of row and column, *τ*_*j*_ represents the random main environmental effect, *ϵ*_*ijrc*_ the plot level residual error modelled with an autoregressive AR(1) x AR(1) variance structure.

#### 2.2.4 Key model outputs

This pipeline produces multiple outputs which can provide valuable insight into the nature of GEI in the data set. Genomic best linear unbiased predictors (GBLUPs) can be calculated for each environment included in the model, as is common in prediction-based modelling of GEI. These GBLUPs provide insight into performance and can be used to derive measures of stability across environments.

Using equation 4, GBLUPs for yield for all varieties were predicted as a linear combination of effects. The GBLUPs in a single environment 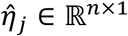calculated in matrix notation as:

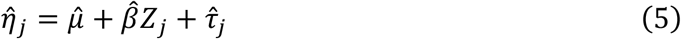

where *τ*_*j*_ is the scalar for each environment. 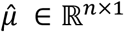 is the vector fixed genotype effects *μ*_*i*_, *Z*_*j*_ ∈ ℝ^*h*×1^ is a vector with elements *z*_*j*1_, *z*_*j*2_ … , *z*_*j*6_ of synthetic environmental covariates for environment *j* and 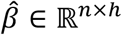 is a matrix of the associated slopes as calculated from the unstructured interaction term between the synthetic environmental covariates and the genotypes.

The nature of the synthetic covariate model allows for of the effect of a single synthetic covariate on the breeding value to be calculated, rather than their collective impact on the GEI (Eq. 5). In this case study this represents the impact of environmental conditions at a single developmental stage can be explored. The deviation/effect for each genotype 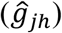 specific to one synthetic covariate (*zh*) can be calculated:

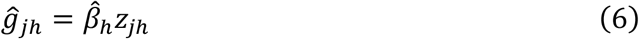

where only the portions of the matrices *Z* and 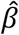 that pertain to synthetic covariate *h* and the associated interactions are utilised.

To determine the effectiveness of the model at capturing the GEI variance, the variance attributed to the synthetic covariates GEI term is calculated as:

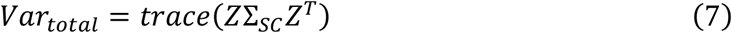

where Σ_*SC*_ is unstructured variance-covariance matrix between the environments from the synthetic covariate model. This model (Eq. 7) does not represent a true value of the genetic variance captured by the synthetic covariates as the model contains a fixed genotype effect it does however represent the GEI variance captured by the model.

Whilst the variance captured by all the synthetic covariates together (Eq. 7) provides insight into model fit and appropriateness, the variance attributed to a single synthetic covariate (e.g. Flowering) provides information on the developmental windows’ relative importance for yield stability. Synthetic covariates contributing small amounts of GEI variance relative to others, in this study, represent time points at which environmental conditions are having less influence on yield. To explore the GEI variance captured by a single synthetic covariate (h) the model can be calculated as per equation 8 (Denis *et al*., 1997; Tolhurst *et al*., 2022).

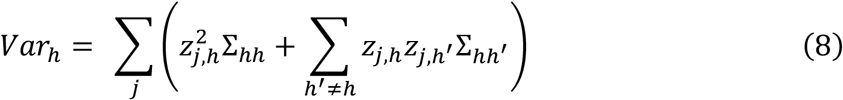

*z*_*j,h*_ is synthetic covariate *h* in environment *j*. Σ_*hh*_ represents the diagonal element of Σ_*SC*_ and 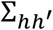 the off diagonal element. Following the Shapley contribution approach half the relevant covariances are included in the contribution of variance by *h* where the covariances are shared between the synthetic covariance of interest *h* and any non-*h* synthetic covariates *(h*^′^).

Further variance decomposition allows for the relative contribution of GEI variance for each environmental covariate to each synthetic covariate. As such within a developmental stage the contribution of each environmental covariate to the GEI variance can be explored. This provides insight into valuable developmental window-environmental covariate combinations which drive yield GEI in this data set. Given the variance-covariance matrix for a specific environmental covariate model is 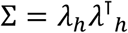 the contribution of an individual environmental covariate (*m*) with in a specific synthetic covariate (*h*) can be calculated in the same fashion as equation 8 utilising the environmental covariate (*m*) in place of the synthetic covariates.

The simplified version of the equation is:

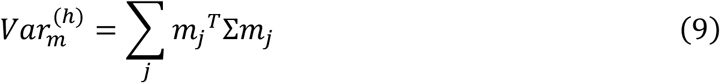

### 2.3 Post Pipeline Analysis

The above pipeline outputs represent the key valuable information that can be extracted from this modelling approach. Below follows further analysis which was conducted to determine more information specifically around yield stability within the context of this case study.

#### 2.3.1 Approaches to understand environmental drivers and stability

Understanding stability and variability with in a well defined TPE is vital to breeding as such the genetic correlations between the environments provide insights into the relationships between these environments as modelled with synthetic environmental covariates.

Genetic correlations between environments due to the synthetic covariates were calculated as per:

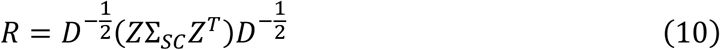

where *D* is the diagonal of Σ_*SC*_. The contributions of synthetic covariates to the genotype by environment interactions for yield were calculated by equation 8 and these results were further partitioned into environment-specific contributions of synthetic covariates by simply calculating the contribution for only a single environment.

From this model performance values and deviations from mean genotypic performance can be calculated for a range of combinations at increasing complexity. For example, the GBLUP across each environment (Eq.5) as well as the GBLUP for each environment at each synthetic covariate (Eq. 6) as such stability can be considered as both stable performance across environments as well as across growth stages and environments. Genotype stability was calculated across environments as root mean squared (RMS) across environments using the deviations from intercepts given equation 6 hence:

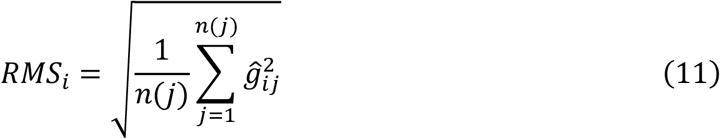

Equation 11 was also utilised to explore stability across the environments at a single developmental window where 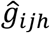 (as per Eq. 6) was utilised in place of 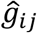.

To create a single measurement of stability across developmental windows and environments was then derived from the RMS values as calculated for each growth stage (h).

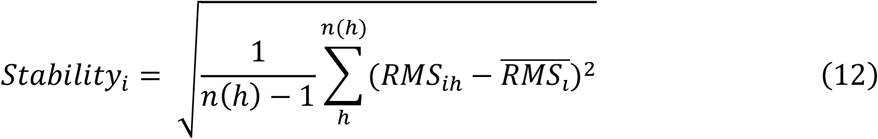

#### 2.3.2. Haplotype stability analysis

Haplotype analysis was conducted using local genomic estimated breeding value methodology to calculate a localised breeding value for a specific haplotype (Voss-Fels *et al*., 2019; Shaffer *et al*., 2026). Haploblocks were defined as SNPs in linkage disequilibrium with a threshold of 0.6 and a tolerance of 3. Tolerance refers to the maximum number of markers not meeting the LD threshold that can be included with in a haploblock. From a total of 6599 SNP markers, a total of 1741 haploblocks were defined across the 7 chromosomes; the haploblocks are the same as those described in Brunner et al. (2026).

Marker effects (*α*) were calculated for each environment using a back-solving approach from the GBLUP with in each environment as per equation 13 then summed based on their assigned haploblock to generate haplotype effects.

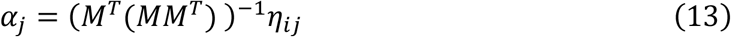

Where M is the centred genotype marker matrix (where the reference allele as 2, alternate allele as 0 and heterozygous as 1 with dimensions genotypes by markers) and *η*_*ij*_ is the matrix of GBLUPs for environment (*j*). Haplotype effects were calculated per environment for the GBLUPs obtained from equations 5 and 6 providing haplotype effects across all synthetic covariates and for each synthetic covariate respectively.

Stability of haplotypes across developmental windows within environments was calculated as per equation 11 where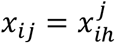was the haplotype effect for genotype i at growth stage h given environment j. To identify the stable blocks of breeding relevance all blocks with perfect stability (RMS=0) were filtered out of downstream analysis due to small to 0 effect on yield. Haploblocks of interest were identified as those with a large range of effects on the trait and chosen for further analysis. Haplotypes at these haploblocks with highest relative positive effects for yield were deemed the top haplotypes.

## 3 Results

The GEI nature of this data set has been described thoroughly in Brunner et al. (2026). The genotypes revealed a moderate level of population structure (PC1 and PC2 account for ∼29% of variance captured) representative of the breeding program nature of the germplasm in this study (Supplementary Figure 1A). The 12 environments represent a range of yield scenarios with site mean yield ranging from 2.5 t/ha in 20MRA to 5.5 t/ha in 21SCW (Supplementary Figure 1B).

### 3.1. Synthetic covariate patterns reveal developmental windows

Flowering date of Spartacus CL was determined for each trial and used as a reference point for developmental windows designated as 0°Cd. Trials did not share identical thermal time lengths and the total shared range of thermal time from sowing to harvest was -1100°Cd to 1000°Cd as calculated from their sowing and harvest dates (Supplementary Table 1). The environmental covariate model was fitted to each of the 11 windows defined with in this range (Figure 2, top) and positive correlations between *λ* (Eq. 3) indicated shared response to these environmental variables (Figure 2, middle).

**Figure 2.**
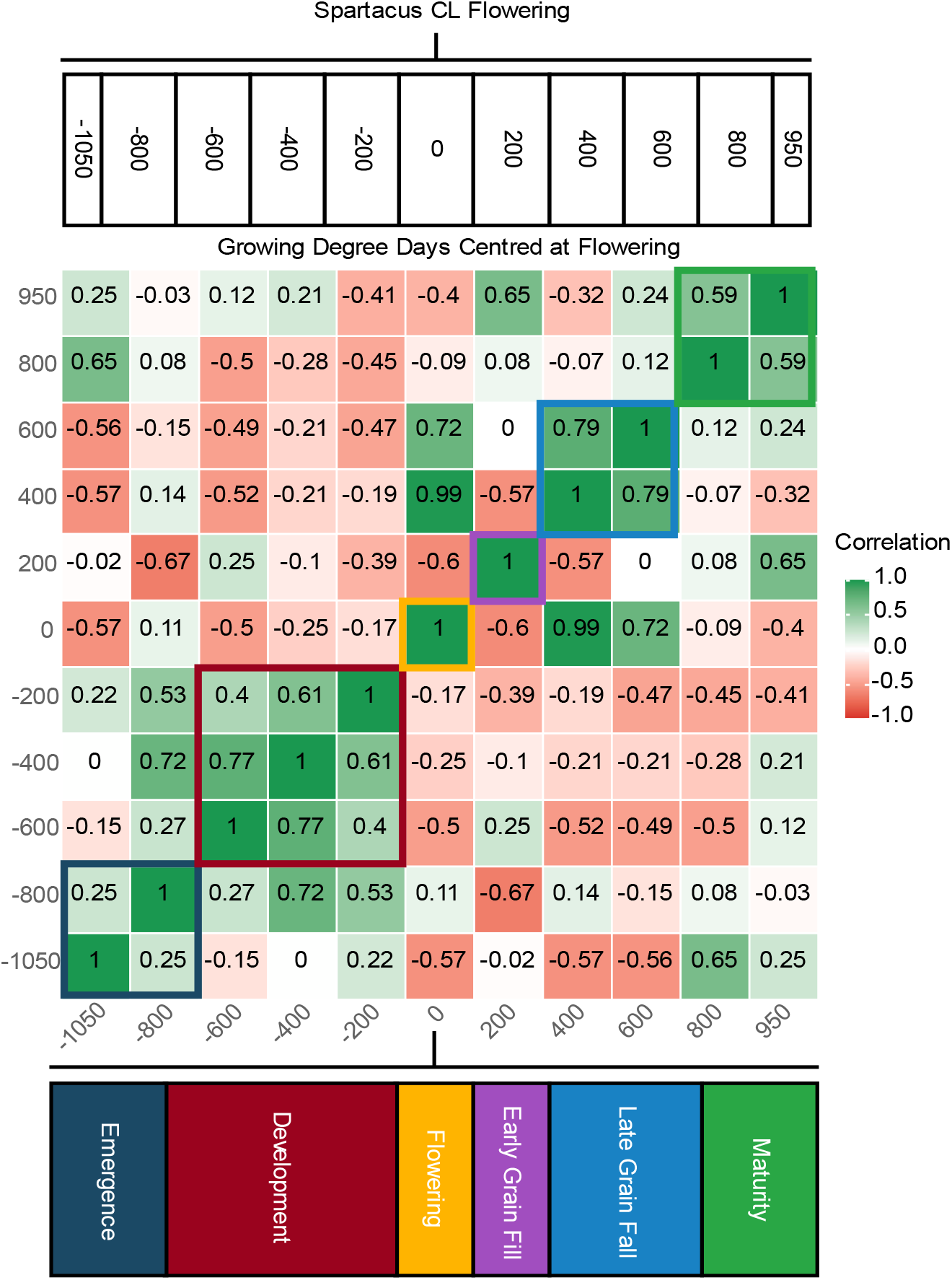
Developmental windows and correlation of their weighted environmental covariate. At the top are presented the 11 initial windows used to identify patterns in λ response to environmental covariates across time. Below is a heat map of the correlations between λ over time between 11 windows for the 12 studied trials. At the bottom are presented the developmental windows chosen to calculate synthetic environment covariates.

Positive correlations between *λ* were observed in blocks with the time from -700°Cd to - 100°Cd before Spartacus CL flowering (Development) being the largest developmental window defined with consistent adjacent positive correlations. Both flowering and early grain fill had no adjacent positive *λ* correlations and hence were kept as individual 200°Cd blocks. Environmental covariates summed over these developmental windows revealed differential environmental patterns. Traits such as rainfall varied greatly across different environments within the same growth stage (24.8mm in 22DND to 186.5mm in 21DND during development stage). Summed minimum temperature also varied greatly especially during late grain fill (148°C in 22YRK to 253°C in 22DND), these variables capture the range of environmental backgrounds utilised in this study (Supplementary Figure 2).

### 3.2. Timing of Environmental Covariates influences yield effect

Yield GBLUPs across environments revealed genotype-specific patterns of GEI (Figure 3A), which can be decomposed to show how environmental covariates at particular growth stages, such as flowering (Figure 3B) and early grain fill (Figure 3C), drive genotypic deviations from the fixed intercepts. Varieties Spartacus CL and Rosalind shared similar GEI profiles with their deviations from the mean effect being almost identical at both flowering and early grain fill (Figure 3B & 3C). Variety RGT Planet exhibited high overall performance, high degree of variability due to weather conditions at flowering but almost no deviations across all 12 environments due to weather at early grain fill. In general, the GEI of the 7 key varieties highlighted appears to have greater deviation at flowering then early grain fill, however the breeding population as a whole show similar variability to that observed for flowering. Notably, parts of the breeding population show positive early grain fill deviations exceeding those of the seven highlighted varieties (Figure 3C), pointing to favourable variation that could be exploited in selection.

**Figure 3.**
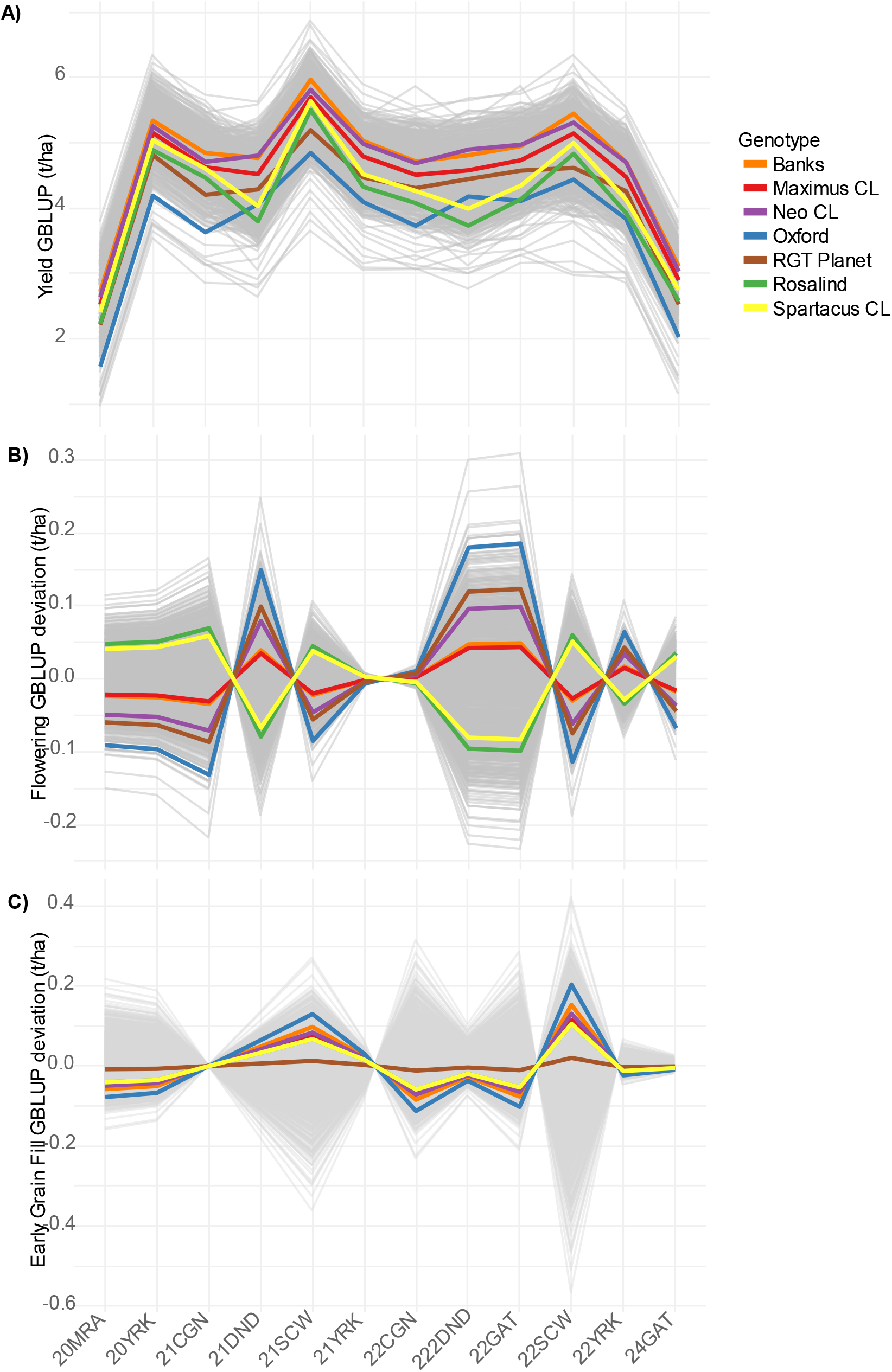
Yield GBLUPs and their deviations from the genotype mean (represented as 0.0) due to synthetic covariates. A) Yield GBLUPs calculated from Eq. 5 incorporating all synthetic covariates. B) Yield GBLUPs calculated from Eq. 6 excluding fixed effects to present deviation from mean due to environmental covariates at flowering. C) As in B but deviations due to environmental covariates at early grain fill.

In this set of 12 environments for barley yield early grain fill is accounting for the greatest percentage of GEI variance with development accounting for the least (Figure 4). The total GEI variance captured by synthetic covariate component of the model was 99.1% based on the other GEI variance being modelled by a diagonal variance-covariance structure assuming no correlation. As this is likely an overestimate the reported GEI variance contributions are all relative and not assumed to be capturing 100% of the total GEI variance present in the data. Environmental covariates at maturity contributed to a large proportion of the GEI variance observed across the synthetic covariates.

**Figure 4.**
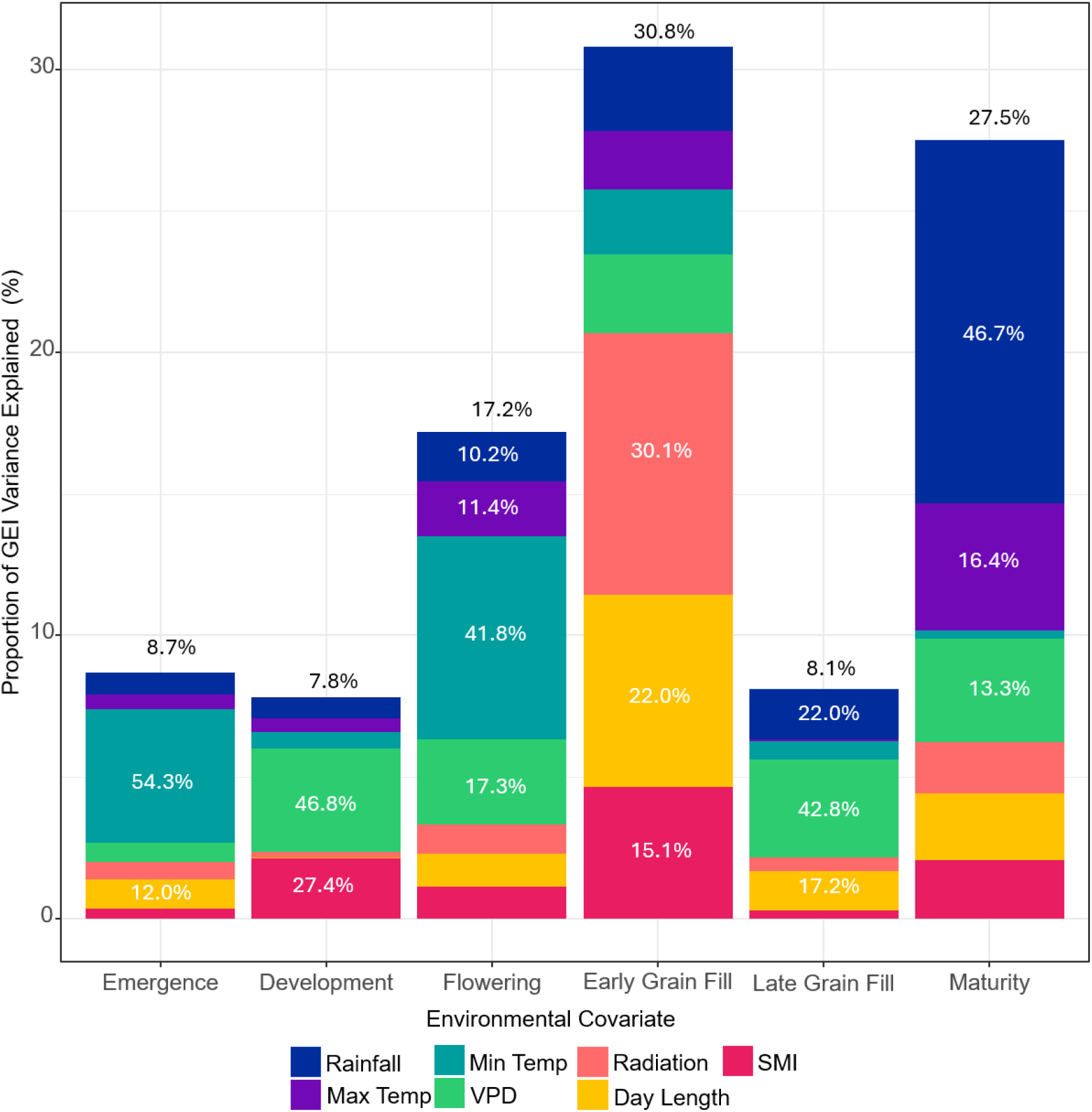
Relative contribution of each growth stage (synthetic covariate) to total genotype by environment interaction captured by the model (percentage shown in black text above individual bars) and relative genotype by environment variance contribution of each environmental covariate for each growth stage (percentage shown in white text within bars).

The specific environmental covariates driving the GEI variance varied greatly across growth windows (Figure 4). No adjacent growth windows demonstrated similar contributions to environmental covariates to GEI variance. Only at flowering and early grain fill were all environmental covariates contributing (Var>6%) to the GEI variance. It is important to note that these variances are conditional hence it can be interpreted that the contribution of minimum temperature at emergence (54.3% of 8.7%) contributes 4.72% of total relative GEI variance. The relative contributions of environmental covariates to GEI variance at specific developmental windows varied across the 12 environments. However, the covariate explaining the least variance was often consistent across environments within a developmental window; for example, radiation consistently contributed the least to GEI variance at emergence (Supplementary Figure 3).

### 3.3. Genotypes can be stable both across and within environments

Further application of this approach to explore GEI patterns across environments observed a high magnitude of genetic correlation between sites with similar profiles of GEI variance attributed to each synthetic covariate (Figure 5). 20YRK and 20MRA which observed similar profiles (Figure 5A) had genetic correlations of 0.99 whilst 21DND and 21CGN which also had similar contributions had a genetic correlation of -0.95 (Figure 5B). Not all environments contributed equally to the genotype by environment variance with 22SCW contributing the most (∼25%) and 21YRK the least (<1%) (Supplementary Table 2). Analysis of the yield variability of varieties performance across environments revealed two of the seven key varieties, Maximus CL and Banks, which had below average variability, hence above average stability, across environments and above average yield GBLUP from the panel (Figure 6A). In contrast Oxford was both very unstable and had a below average GBLUP for yield across environments. Stability across developmental windows revealed that Oxford in general is an unstable variety, however, is found to be highly stable across environments at late grain fill (Figure 6C). RGT Planet was unstable at the early growth stages but among the most stable at maturity and early grain fill, the two time points are contributing the most to GEI variance for yield. Comparing variability across environments at flowering and early grain fill found that Maximus CL is the only variety of the 7 to have above average stability at both developmental windows (Figure 6B), however generally across all windows those varieties that are stable across environments are so at more than one growth stage (Supplementary Figure 4).

**Figure 5.**
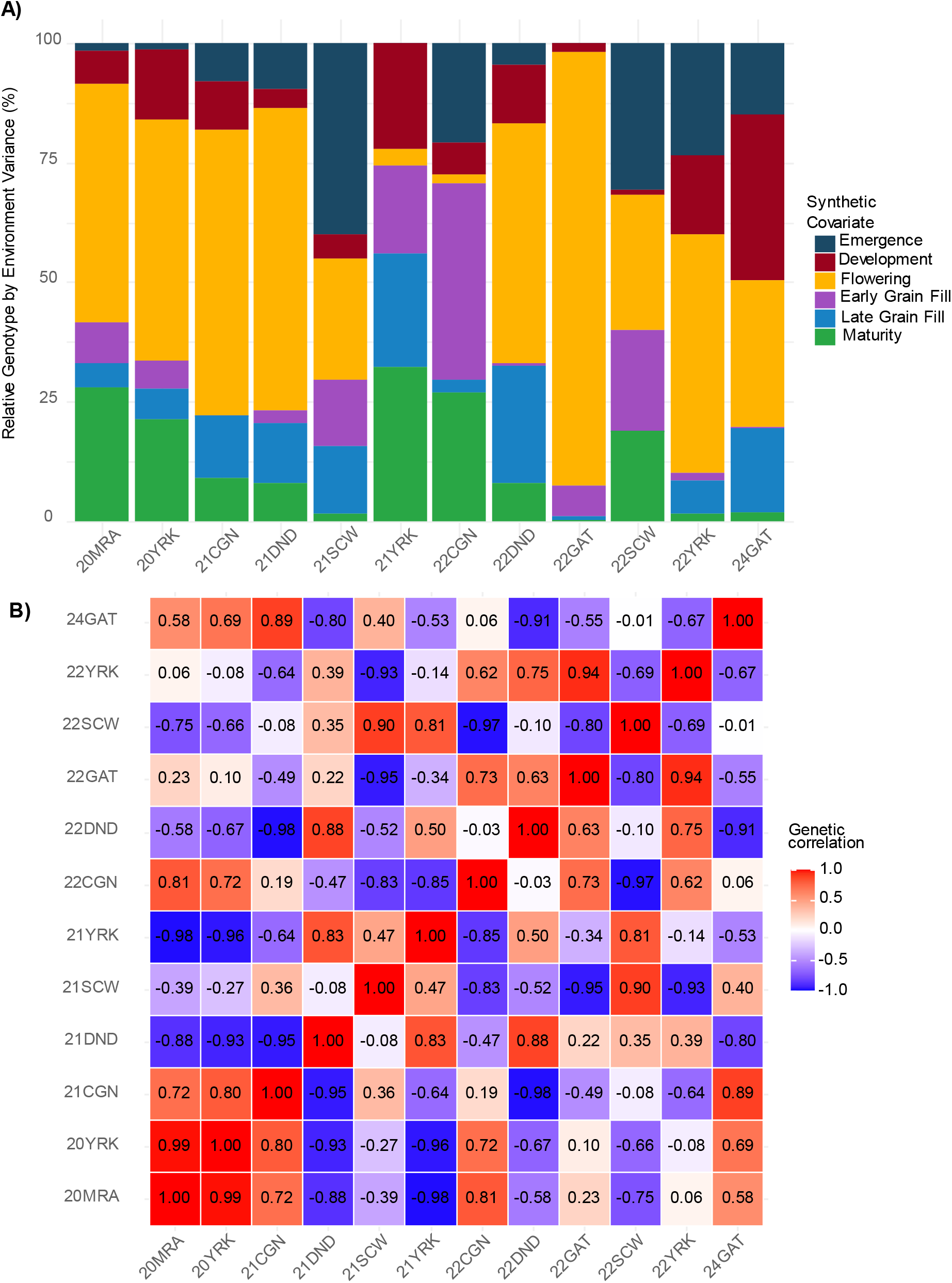
Yield variance and shared variance across environments. A) Environment specific contributions of synthetic covariates to relative genotype by environment variance (%). B) Pair-wise genetic correlations between environments, calculated as per equation 10.

**Figure 6.**
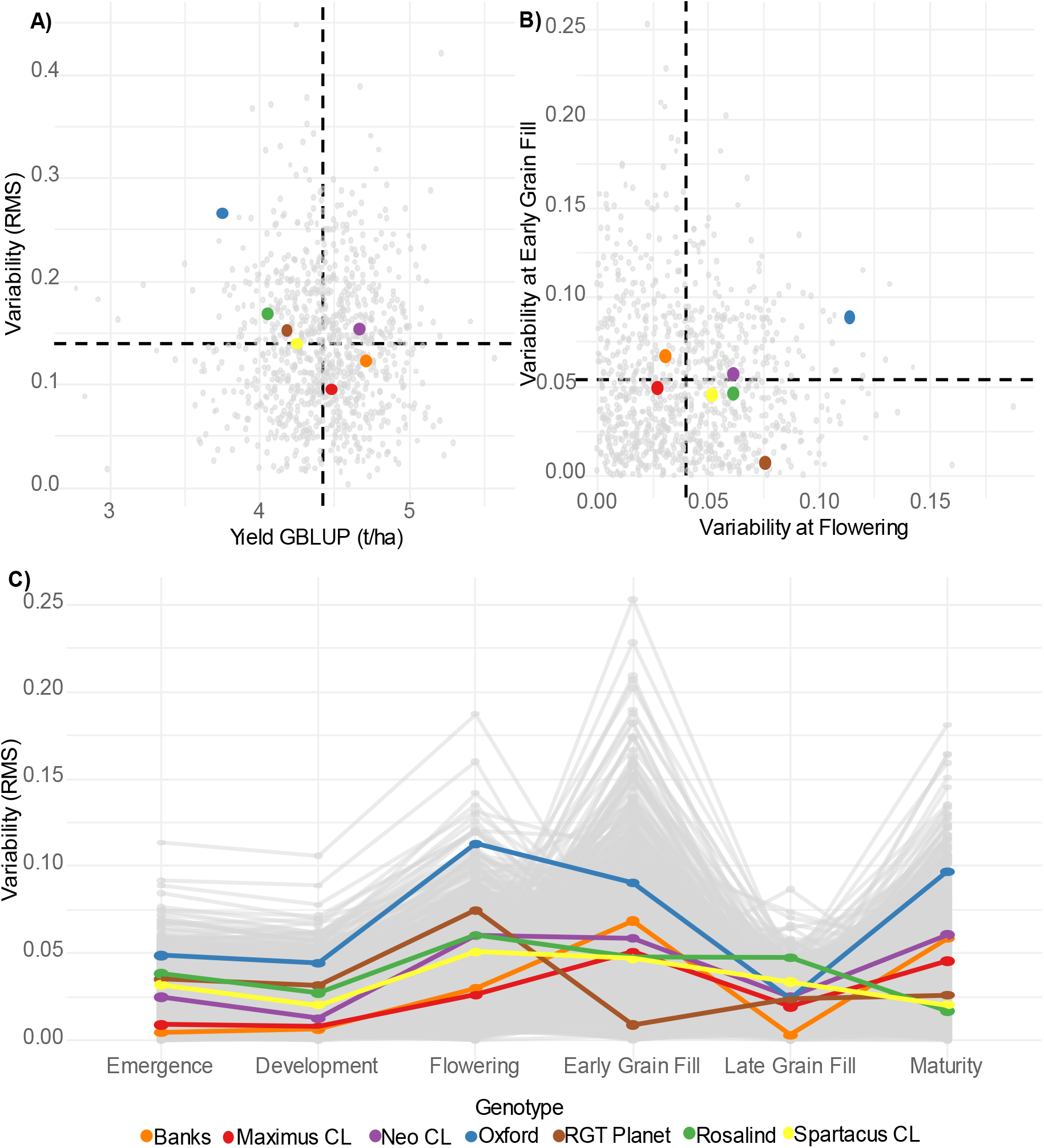
Yield variability and performance across environments. A) Variability across environments (RMS as per Eq. 11) against Yield GBLUP, vertical dashed line represents mean yield, and horizontal dashed line represents mean stability. B) Variability at flowering and early grain fill. C) Variability across environments and stability across growth stages (as per Eq 12). Key genotypes coloured as per legend.

Haplotype based stability revealed that generally stable and unstable haplotypes are shared across multiple environments with correlations for haplotype stability ranging from -0.12 to 1 (Figure 7A). Stability across growth stages was moderately conserved in 21CGN and 22GAT with a correlation of 0.82 (Figure 7B). Despite the correlation the two most unstable blocks b000304 and b001604, previously identified as haploblocks with GEI in Brunner et al. (2026), had contrasting top haplotypes. The top haplotypes in 21CGN for block b001604 and the top haplotypes in 22GAT for b000304 represented the least stable haplotypes across both environments. Notably, the highest performing haplotypes withinb001604 in 22GAT also exhibited the greatest stability. This pattern was conserved when contrasting two environments with less correlated haplotype stability 22GAT and 24GAT (r=0.32; Figure 7C), though across all the haplotypes a more fanned distribution was observed indicative of haplotypes stable in one environment but unstable in the other.

**Figure 7.**
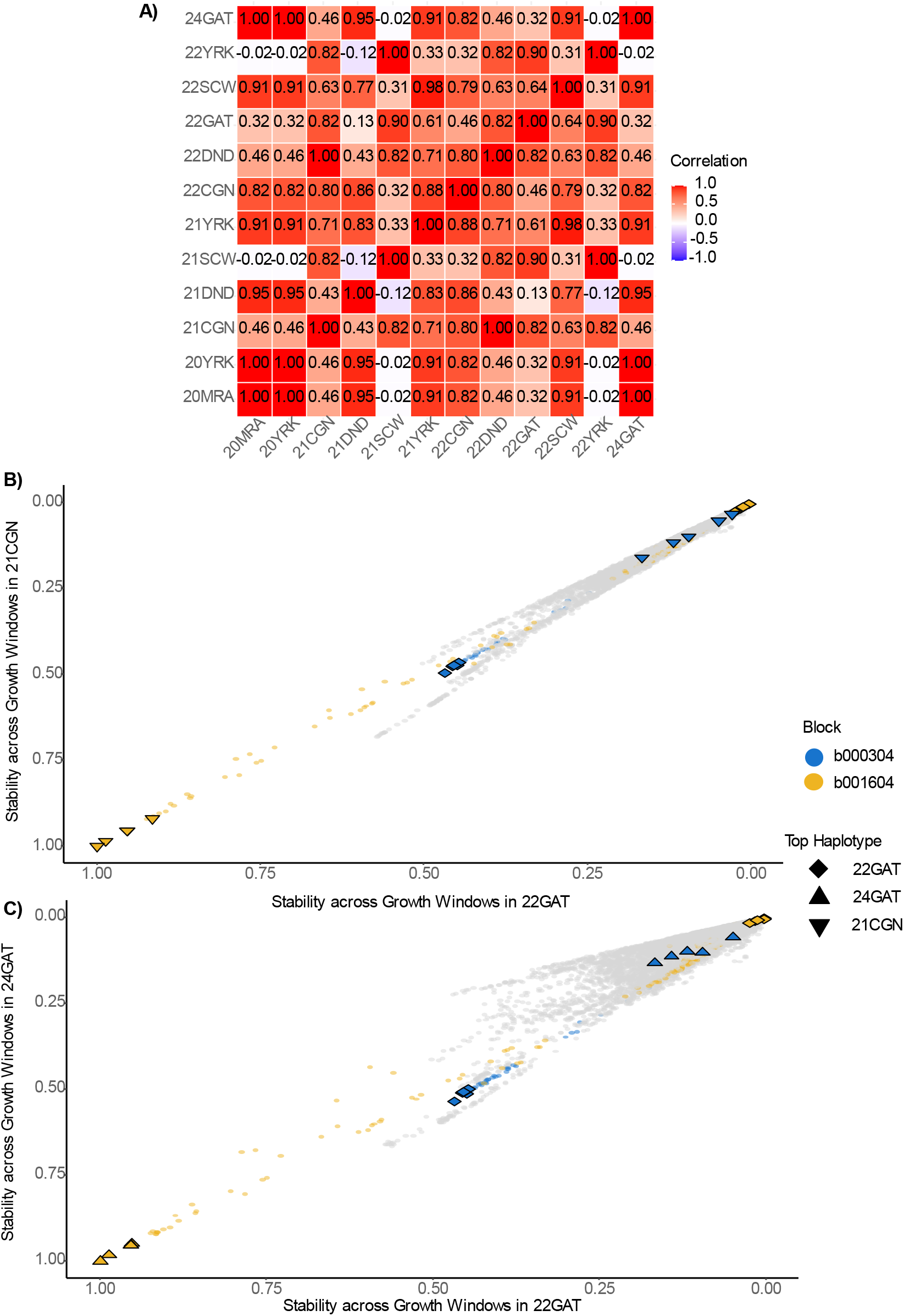
A) Correlation of haplotype stability across growth stages between each environment calculated as per Eq. 11. B) Stability across growth stages of haplotypes in 22GAT and 21CGN. C) Stability across growth stages of haplotypes in 22GAT and 24GAT. Blocks b000304 and b001604 coloured in blue and yellow respectively. Top 5 haplotypes in each environment for b000304 and b001604 denoted by diamonds and triangles.

## 4 Discussion

### 4.1. Synthetic covariate modelling reveals developmental window specific patterns of GEI for barley yield

In this study, we aimed to dissect GEI at the level of individual growth stages within environments, before examining how these patterns manifest across environments, which is an approach that offers considerable opportunity for improving our understanding of the environmental drivers of yield GEI. Applying our synthetic environmental covariate pipeline to this dataset revealed patterns of environmental response aligned with known growth stages in barley (Figure 2). Physiological understanding of barley growth stages has historically revealed key developmental transitions that carry distinct environmental sensitivities (Miralles *et al*., 2021). Flowering time, for example, has been shown to represent a narrow period during which barley is particularly sensitive to environmental conditions and when stresses frequently occur (Dreccer *et al*., 2018; Bratković *et al*., 2024; Heidariask *et al*., 2026), whilst the longer vegetative stage during development has been found to be less responsive to small environmental shifts. Changes in environmental response of genotypes is often considered across environments as GEI however within a single trial the environment does not remain static nor does the response which is specific to a growth stage. Whilst the plants genotype does not change across growth stages gene expression alters throughout a plants life (Liew *et al*., 2020). Contrasting responses were observed to environmental conditions at different time points (Figure 3B and 3C) suggesting that the GEI at a specific growth stage-by-environment combination contribute unique and distinguishable patterns to the overall yield GEI (Wiegmann *et al*., 2019). These growth stage-specific patterns of GEI, once characterised within environments, provide a biologically interpretable basis for understanding the broader GEI signal observed across environments, and therefore a basis for strategic design of crop improvement programs.

In this data set it was identified that the environmental conditions at early grain fill contributed the most to the overall GEI variance for barley yield. This is aligned with expectations, as this stage of development contributes significantly to grain weight (Becheran *et al*., 2022; Ullah *et al*., 2026). This aligns further with radiation at early grain fill accounted for ∼10% of GEI variance from the synthetic covariate model. Increased radiation post-anthesis in cereals increases photosynthesis and final yield (Murchie *et al*., 2002; Zhang *et al*., 2022). Environmental conditions at maturity contributed the next greatest amount to overall variance. This pattern is less intuitive as it can be assumed that photosynthesis and nutrient remobilisation has slowed or stopped by this time point. However, ∼47% of this variance is attributed to rainfall. It is likely that rather than a direct physiological response being captured this is indicative of a trial management limitation, as high rainfall can delay harvest or cause yield losses through lodging (Bolland, 1984; Laidig *et al*., 2022). At all growth stages, except maturity, minimum temperature contributed more than maximum temperature to GEI variance. As environmental data is on a daily scale the minimum temperature is representative of the night temperature which has been found to contribute greatly to yield variability in wheat (Schulthess *et al*., 2026, Preprint). Despite these generalisations that are in line with physiological understanding of the complex landscape of barley yield it is important to note that these contributions to the variance are generally environment specific (Supplementary Figure 3) and thus depend on the dataset analysis. In particular, the 12-trial dataset did not capture all possible stresses within the broader TPE, such as substantial pre-flowering drought or heat stress, which were limited or absent in the studied trials (Heidariask *et al*., 2026). Given that these contributions are largely environment-specific underscores the value of examining GEI at the resolution of individual growth stages within environments, whereby environments that appear broadly similar in overall performance may differ considerably in the specific developmental windows and environmental factors driving yield. This developmental window-level perspective not only provides a biologically interpretable basis for environmental grouping and TPE definition, but more broadly demonstrates that dissecting GEI beyond the whole-season level has the potential to substantially improve our ability to characterise and exploit genotype-specific environmental responses in breeding. The results of this case study are consistent with previous research, despite the relatively small number of environments evaluated. Applying this approach to larger breeding program datasets could enable a more detailed dissection of disentangle GEI than has previously been possible.

### 4.2. Synthetic covariate modelling potential, extensions and limitations

The primary aim of this paper is to present the synthetic environmental covariate pipeline as a new approach to dissect GEI in complex data sets at scale, while exploring opportunities for extension in breeding and research application. These include haplotype by environmental covariate interactions or designed multi-environment trials to answer physiological questions on a large scale. Therefore, it is important to stress the flexibility of this approach. The case study presented for barley yield presents one such way to analyse GEI across environmental covariates, growth stages and environments however it is not the definitive manner by which this pipeline can be applied and is also limited by the number of locations in the dataset.

The synthetic covariate model fit in this study appears to capture ∼99% of the GEI variance present in the dataset when contrasted with the residual GEI interaction between genotypes and environments. The computational restrictions preventing the inclusion of more complex variance-covariance structures to the GRM by environment main effect (i.e. the environment effect not captured by the synthetic covariates) pose a limitation to this analysis. This limitation could likely be bypassed with a reduced number of genotypes or environments. In this instance it does not pose a direct impact on the results as the additive GEI due to environmental covariates is the term which this pipeline is designed to explore.

The initial environmental covariates utilised in this proof-of-concept study were restricted to summed daily values. Further work should investigate the inclusion of more physiologically meaningful variables that better capture important specific stresses, such as frost, heat, and drought. Crop modelling and remote sensing approaches could be particularly valuable for deriving such variables (Chenu *et al*., 2011; Potgieter *et al*., 2021). For example, this study considered only rainfall and a soil moisture index (SMI) estimated from generic physical soil properties and climatic information, whereas drought stress depends not only on soil and climatic conditions but also on crop status, including root distribution and transpiring leaf area, which can vary substantially among environments (Chenu, 2015; Vadez *et al*., 2024).

Similarly, rather than relying on maximum temperature alone, recent work in wheat has shown that the number of hours above a critical temperature threshold can better explain yield responses (Ullah *et al*., 2024). In addition, with advancements in sensing and remote sensing the amount and accuracy of environmental data that can be collected in a single trial is growing rapidly (Zhao *et al*., 2025; Fernandez *et al*., 2026). Resources such as the NASA POWER database provides hundreds of climate parameters for any given location on any given day though with less accuracy then in field weather stations (Zhang *et al*., 2007). As such the environmental covariates utilised are limited only to computational and statistical capacity of dataset and the model. This presents an alternative application of the pipeline for feature selection to determine which environmental variables contribute the greatest amount of GEI variance for the trait of interest. In such applications the pipeline could be used to either remove those environmental covariates with minimal contributions or retain those that capture the most variance (Clouvel *et al*., 2025). Overall, this model presents a rare opportunity to assess the contribution of such environmental variables in the same model framework as the final model, utilising the same inputs and overall model structure, effectively minimising additional unnecessary error.

Synthetic environmental covariates are derived from groupings of these environmental covariates. In the case of this study these groups were defined based on developmental windows, to explore the temporal change in GEI. However, the same set of environmental covariates could have been grouped any number of ways. An alternative approach would be to derive synthetic covariates based on categories of weather variables, for example, combining all rainfall measurements across the growing season into a single synthetic covariate, rather than partitioning environmental data by growth stage. This approach would limit the ability to explore the temporal dimensions of GEI but would provide greater ability to interpret the response to specific correlated yet distinct environmental characteristics align to a breeding objective or research question.

Given their interconnected nature the combination of potential environmental covariates and their groupings into synthetic covariates rapidly becomes a large number of possibilities.

Exploration of these combinations using genetic algorithms or machine learning approaches presents an opportunity for future research (Kusmec *et al*., 2024; Souaibou *et al*., 2025).

However, in many cases would be unnecessary as the flexible nature of this pipeline allows for the development of synthetic covariates to answer specific questions for both researchers and breeders.

### 4.3. Breeding applications of approach

Stability is a key breeding target for yield, quality, disease resistance and other traits, and is commonly defined as consistent performance across environments. However, stability within an environment is less often considered, despite both environmental conditions and genotypic responses changing across the growing season. This pipeline adds a developmental dimension by quantifying the consistency of genotypic performance across changing conditions within individual growth-stage windows. Rather than selecting only for broad stability, breeders could therefore target stability, or favourable responsiveness, within the developmental windows contributing most to yield GEI. For example, if early grain filling is the most influential window, selection could favour genotypes that are stable or respond positively during this period. Variation attributable to within-season responses was smaller than variation among environments, suggesting that selection pressure on this component may not need to be as stringent (Figure 6). The proportion of variance attributed to each growth stage could also identify time points where response to environmental covariates is most important. RGT Planet provides an example, showing stability across environments during early grain fill despite lower stability when all developmental windows were combined (Figure 3). Given the relationship between weather during early grain fill and yield GEI, together with the importance of this stage for grain weight, this developmental window represents a potential stability target. Similarly, synthetic covariates derived by weather category rather than growth stage could be used to assess varietal stability across specific weather patterns.

Stable genotypes may nevertheless carry unstable haplotypes, and vice versa. Haplotype analysis can therefore identify genomic regions with stable or large effects across synthetic covariates. Correlations in haplotype stability between pairs of environments revealed strong haplotype-level GEI (Figure 7A), while comparisons across growth stages illustrated trade-offs between stability and effect (Figure 7B and 7D). In both 24GAT and 21CGN, the most beneficial haplotypes for yield at block b001604 were also the most unstable, whereas the best-performing b001604 haplotypes in 22GAT were the most stable. Such information could help breeding programs interpret the relationship between stability and effect at specific chromosomal segments within particular environment types (Brunner *et al*., 2026).

For stability to be relevant to breeding, the environments used must represent the TPE (Cooper *et al*., 2023). This is primarily achieved through experimental design but can also be supported by environmental clustering, which aims to group environments while minimising re-ranking of variety performance between groups (Russell *et al*., 2003). Clustering has been based on genetic correlations, latent factor responses, environmental variables or geography (Russell *et al*., 2003; Smith *et al*., 2021; Bustos-Korts *et al*., 2022; Elmerich *et al*., 2023).

Although environmental clustering was not applied here, the pipeline provides potential criteria for doing so. Responses to specific synthetic covariates (Figure 3B and 3C) reveal environments with differing levels of crossover GEI, allowing clustering based on GEI patterns at key time points or at the growth stage contributing most to GEI variance. Because these patterns varied across growth stages in this dataset, such clusters would be stage-specific rather than generalisable across the season.

Environments could also be clustered according to the proportion of variance attributed to each synthetic covariate. This showed moderate agreement with genetic correlations; for example, 20MRA, 20YRK and 21CGN had similar contributions from flowering, late grain fill and development and were highly genetically correlated (Figure 5). However, similar variance profiles did not necessarily imply the same correlation direction, as 21DND had comparable variance contributions but was strongly negatively correlated with the other three sites. Self-organising maps provide another potential clustering approach A limitation of clustering directly on environmental data is that clusters may primarily reflect the environmental main effect and overall productivity rather than genotypic response (Lacaze and Roumet, 2004; Malosetti *et al*., 2013). In contrast, synthetic covariates weight environmental variables by their importance to trait GEI, potentially providing clusters that better reflect genotype-specific environmental responses and are partially predictable for use in breeding.

The model can also be extended to prediction in new environments when relevant environmental covariates are available (2023). With environment fitted as a random term, performance in new environments can be predicted together with its prediction error. Model parameterisation should therefore reflect the research question. When the aim is to interpret or extrapolate covariate loadings, or predict into future environments, the population-average synthetic-covariate slope must be included as a fixed effect rather than absorbed by the random environment effect (Piepho and Blancon, 2023; Piepho and Williams, 2024). This provides an alternative to approaches based on correlations between latent factors and environmental variables (Fradgley *et al*., 2025) or latent factors in combined with environmental covariates (Tolhurst *et al*., 2022). Future research should evaluate the ability of this model to predict genotype performance in unseen environments using observed or projected environmental covariates. Projected weather conditions could also be used to explore varietal performance under future climate scenarios.

### 4.4. Conclusions

As environmental conditions become more variable, understanding GEI is increasingly important for field-trial interpretation and breeding decisions. Environmental responses also change within environments as crops develop. By combining quantitative genetics, environmental characterisation and crop development, our approach identifies when during development environmental conditions contribute to yield variation across environments.

Although it does not provide the mechanistic detail of dedicated physiological experiments or crop-growth models, synthetic environmental covariates provide a more explicit link between GEI and measured environmental conditions than purely latent models. The approach also revealed that yield stability varies across development, although highly unstable varieties tended to remain unstable across stages. This temporal decomposition identifies stage-specific stability as a potential breeding target. Trade-offs between effect magnitude and stability were also evident among high-performing genotypes and large-effect haplotypes.

Together, these results demonstrate how synthetic environmental covariates can extend Finlay–Wilkinson regression beyond describing GEI to identifying the developmental periods and environmental conditions associated with genotype responses, providing breeders with a more biologically interpretable framework for understanding and selecting for yield stability.

## Abbreviations

(BLUE): Best Linear Unbiased Estimators
(FA): Factor Analytic
(GBLUP): Genomic Best linear Unbiased Predictors
(GDD): Growing Degree Days
(GEI): Genotype by Environment Interactions
(GRM): Genomic Relationship Matrix
(LD): Linkage Disequilibrium
(Local GEBV): Local Genomic Estimated Breeding Value
(RMS): Root Mean Squared
(SMI): Soil Moisture Index
(SNP): Single Nucleotide Polymorphism
(TPE): Target Population of Environments
(VPD): Vapour Pressure Deficit

## 5.1 Acknowledgements

The authors thank the technical and breeding staff at InterGrain Pty. Ltd for providing the germplasm and genotypic data used in this study.

## 5.2 Author Contributions

Conceptualisation – SMB, HR, KVF, LH; Data Curation – SMB, DG, SB, DM, HR; Formal Analysis – SMB, BH; Funding Acquisition – LH, KVF; Investigation – SMB, KC; Methodology – SMB, HPP, HR; Project Administration – LH; Resources – DG, SB, DM, LH, KVF. Supervision – LH, KVF, HR; Visualization – SMB; Writing – Original Draft – SMB; Writing – Review and Editing – SMB, HPP, KC, BH, DG, SB, DM, LH, KVF, HR.

## 5.4 Data Availability

The data that support the findings of this study are available from InterGrain Pty Ltd, but restrictions apply to the availability of these data, which were used under license for the current study and so are not publicly available. Data is, however, available from the authors upon reasonable request and with permission of InterGrain Pty Ltd.

